# FlowFI: an interactive graphical software package for bespoke design of imaging parameters in flow cytometry to explore morphological diversity in bone marrow megakaryocytes

**DOI:** 10.64898/2026.05.19.725920

**Authors:** James B. Wilsenach, Sonia Fonseca, Sebastian E. Ahnert, Edyta E. Wojtowicz

## Abstract

**Background:** Imaging flow cytometry (IFC) provides a high quantity of single-cell morphological data, yet the field lacks open access tools for designing interpretable, bespoke parameters. In particular, rare and atypical cell populations where well annotated data is limited, are negatively affected.

**Results:** We present Flow cytometry Feature Importance (FlowFI), an open-source graphical software for bespoke image parameter design and analysis. FlowFI provides a suite of image parameter options combining data across multiple channels and markers, tailored digital noise reduction (reducing noise resulting from common flow cytometry ultra-high image acquisition modalities), and a scalable, unsupervised feature selection pipeline that allows experimentalists to refine image-derived parameters iteratively, with a novel ensemble subsampling approach that provides robust feature importance scoring. We validated FlowFI using data from a rare and heterogenous bone marrow cell type, megakaryocytes, demonstrating that the tool can successfully identify novel, discriminatory morphological features to improve the purity of selected cell populations and gating strategy.

**Conclusion:** FlowFI’s core functionalities are interacted with through an intuitive user interface for researchers with options to export data directly to common image and flow cytometry software formats. With this in mind, FlowFI offers a scalable way to both feature design, and feature refinement using a range of approaches to manifold learning, augmented by a data efficient bootstrap subsampling approach for unsupervised parameter recommendations in the big data regime. The software also introduces a new feature selection measures based on common manifold learning methods in the space inspired by the Uniform Manifold Approximation and Projection (UMAP) algorithm and finds performance comparable to existing methods. FlowFI provides a versatile testing ground for future developments in broad and dynamically developing areas of research including single cell analysis, label-free sorting and intra- and inter-cellular interaction analysis, while ensuring interoperability with current research workflows. Desktop installation options as well as detailed documentation can be found at https://github.com/EarlhamInst/FlowFI

## Background

Flow cytometry revolutionized immunology and haematology by providing both ultra high-throughput and single cell resolution. In conventional flow cytometry each cell (or event) is represented as a single pixel on the screen. In contrast, in Imaging flow cytometry (IFC), each cell is represented by a rich set of spectral features, along with images. These additional modalities make IFC very attractive for use in analysis as well as cell sorting (Image Activated Cell Sorting, IACS) for downstream transcriptional, proteomic or genomic analysis. (1,2). IACS enables high-throughput morphological cell characterisation, cell quality assessment, and, for heterogeneous populations, sample purity refinement (3). While several high-throughput cell-imaging instruments are available (4–6), the integration of morphology recording and sorting of single cells for genomic analysis is currently very limited, with the BD FACSDiscover S8, recognised as the first imaging-enabling full-spectrum cell sorter on the market (7), increasingly adopted in the field (8–10). The S8 produces images reconstructed from electrical pulses emitted by a blue laser split into a set of ‘beamlets’. This is the only system that allows for exporting of raw image files produced (11). This platform provides an array of image-based parameters for cell analysis and sorting and is increasingly focusing on Machine Learning (ML)-enabled biomarker identification (12,13). These methods usually require a large amount of well annotated data. However, for rare or atypical cells with heterogeneous morphologies, this can be challenging to acquire.

In this paper, we explore a case study of a rare, morphologically heterogeneous population of bone marrow cells, megakaryocytes (MKs). These cells constitute less than 0.1% of all bone marrow cells (14) and yet are responsible for the production of millions of platelets every day, while also playing an important role in the immune response (15–17). MKs are characterised by varying amounts of DNA (ranging from 2n up to 128n DNA copies denoted as ploidy), which results in highly variable morphology in relation to their overall size and ratio of the nucleus to the cytoplasm (Fig. 1) This morphological heterogeneity has been linked to distinct MK functions in the BM, primarily producing platelets or providing the support environment for haematopoietic stem cells (18). MK characteristics, including fragility, irregular shape and a tendency to adhere to other cells, make MKs difficult to identify and isolate as single cells using currently available high-throughput methods (19). We use this exploration of bone marrow MKs to illustrate the pressing need for unbiased, bespoke parameter and explicit marker construction to further characterise atypical cells while remaining interpretable and interoperable with existing sorting strategies, analysis pipelines and software (20).

**Figure 1.**
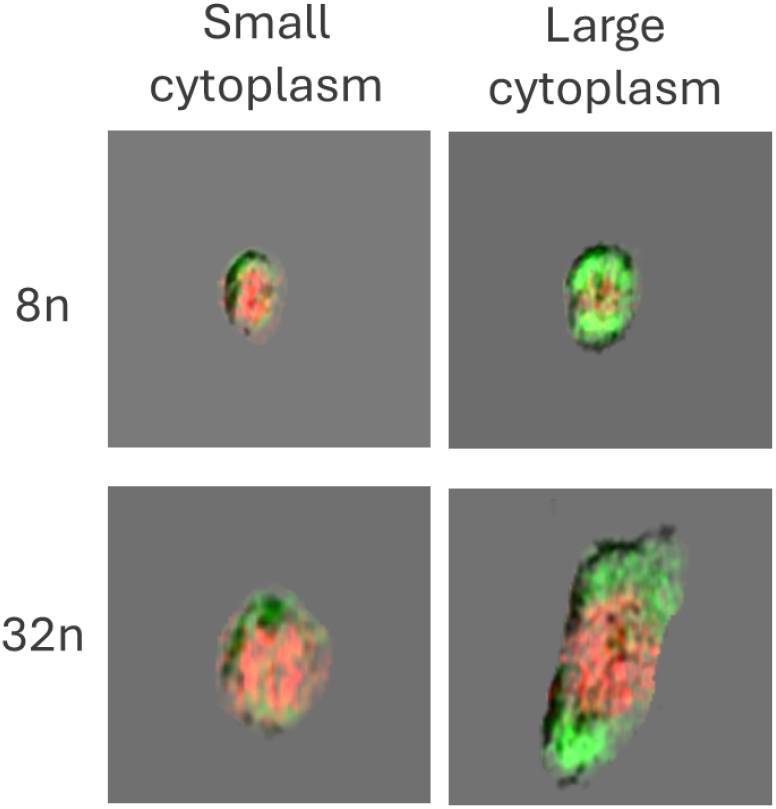
Representative images of megakaryocytes classified according to their ploidy and cytoplasm size: 8n-small, 8n-large, 32n-small, 32n-large. The images were acquired with BD FACSDiscover S8 and analysed in FlowJo v10.10 with BD CellView Lens plugin. The nucleus was stained with DRAQ5 (red) and cytoplasm with CD41-FITC (green).

We present **Flow** cytometry **F**eature **I**mportance (FlowFI), a Graphical User Interface (GUI) enabled software tool for manifold learning-based marker identification, image preprocessing, and design of bespoke parameters specifically designed for images obtained through imaging flow cytometry, though images acquired using other methods can be used as well such as microscopy. FlowFI provides noise reduction steps tailored to reduce the specific anisotropic noise generated during high throughput image acquisition, for example due to the specific noise signatures produced by the BD FACSDiscover S8 which uses waveforms produced through Orthogonal Frequency Domain Multiplexing (OFDM) (see Supplementary Section 1.1) for image reconstruction (21). In addition, FlowFI allows users to design the preprocessing tailored to their own setups and experimental needs, such as by reducing blurring caused during the Time Delay Integration (TDI) step on camera-based imagers (22). FlowFI preprocessing improves upon the signal to noise ratio in multichannel staining and also provides several image segmentation options that allow data from multichannel data to be combined thereby extracting complex multichannel relationships. These preprocessing options both serve as a basis for analysis as well as possible inputs for downstream analysis such as image-based learning.

Modern flow cytometry systems can produce a wide range of parameters. Existing tools provide comprehensive cluster analysis tools that can inform identification of important subpopulations and specific markers allowing their identification within a cell population (23,24). FlowFI uniquely provides an unsupervised feature selection tool for “first look” refinement of new or existing parameters to determine whether novel or existing parameters have the discriminatory power to be useful in downstream analyses. The tool uses ensemble learning (25) combined with bootstrap subsampling (26) to improve scalability of machine learning and conventional feature selection methods to large datasets that can be easily run locally in real time. This tool is designed to complement existing software within the flow cytometry ecosystem as well as FlowFI’s feature design capability, providing feedback on feature usefulness within seconds to minutes (depending primarily on the number of parameters) for iterative parameter improvement.

The following sections provide an overview of FlowFI’s implementation as a user interface and platform for experimentalists to design and refine novel parameters for use in cytometry. In later sections we demonstrate a possible workflow for implementation of the platform to atypical cells and for validation of refined machine learning and image analysis methodologies employed.

## Implementation

FlowFI combines image processing and analysis tools for use within the flow cytometry experimental ecosystem. The specific implementation of FlowFI’s core functions follow two major streams, the creation of new bespoke image parameters with specialised noise preprocessing options designed for the high noise environments of extremely rapid, high throughput image acquisition, and the analysis of these parameters in the context of others to determine relative importance for possible downstream gating or analysis tasks.

### The Design-Refine Loop

FlowFI’s two principal components, the design and refine sweets of features are designed to work in a complementary fashion, possibly involving multiple iterations. Fig. 2 shows the core functionality loop. Starting from acquisition FlowFI allows for visualisation and interactive preprocessing of events in order to improve signal-to-noise ratios, define masks over a specific channel for analysis. The preprocessed image is then quantified into a single summary statistic or parameter using a range of quantification options, these include straightforward counts and complex geometric summaries of the image combining information across multiple channels. The resulting pipeline can be used to batch process a set of images and output a CSV or augment an existing Flow Cytometry Standard (FCS) file to include the new parameter values (27). FlowFI includes options to make these broadly consistent with existing flow cytometry analysis software such as FlowJo (28).

**Figure 2.**
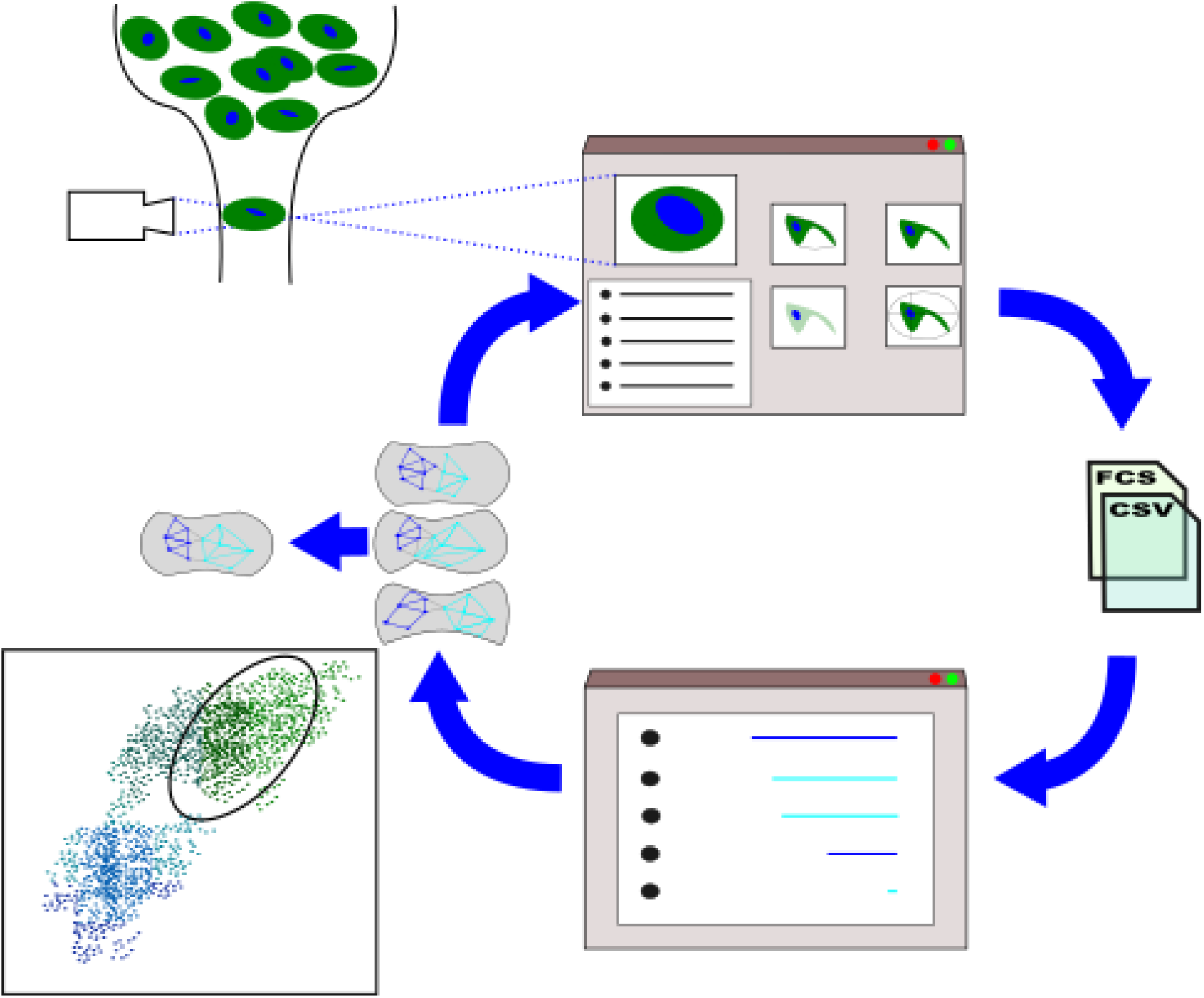
Illustrative overview of the design and refine workflow in FlowFI. High throughput cell morphology data and multichannel images are captured by the imaging flow cytometer. Feature extraction in FlowFI is accomplished through exploring available data and constructing batch processing scripts that first visually preprocess the images and then quantify morphological data according to user specifications, both general and highly specialised multichannel quantifications options are provided. Extracted features can then be compared using the feature importance analysis tool. It uses fast, robust manifold learning algorithms to subsample the data, perform clustering and spectral embedding which are aggregated to rank the existing features. Rank comparison then allows for the further refinement of features in the design tab.

The second component of FlowFI’s core functionality is its suite of refinement features. These are a collection of machine learning-based feature analysis methods that have been optimised and augmented to work efficiently on large flow cytometry datasets (29). The main purpose of these importance measures is to determine which parameters are most likely to yield useful insights. Here, usefulness or importance to the user is known to be highly dependent on the underlying data structure, and user-specific needs e.g. clustering, gating or regression tasks (30–33). Our framework utilises a bespoke subsampled bootstrapping procedure that works across a range of measures to refine the parameter set at scale, providing multiple possible measures that prioritise different structural properties of the data such as globally or locally structure preserving, highly informative or highly representative. In addition, the framework provides empirical confidence intervals for each measurement for statistical rigour (see Supplementary Section 2.1).

The majority of measures available utilise a graph-based approach to manifold learning that determines the most important features (parameters) underpinning the structure of the data space. Graph-based manifold learning provides a robust, non-parametric common framework for analysing unstructured data (34,35). Importance scores from the refinement step can be used to inform the user if the new parameters they produce are potentially useful for analysis and which parameters need to be further improved.

### Feature Design Workflow

FlowFI is a fully visually-enabled user interface for flow cytometry analysis. Fig. 3A shows how feature design is achieved within the programs design tab. FlowFI places the raw image side by side with the preprocessed image for comparison. Specific Quantification options for parameter calculations are available in the *Quantify* submenus. Batch processing is achieved similarly by selecting a folder for batch processing in the *Parameter* menu.

**Figure 3.**
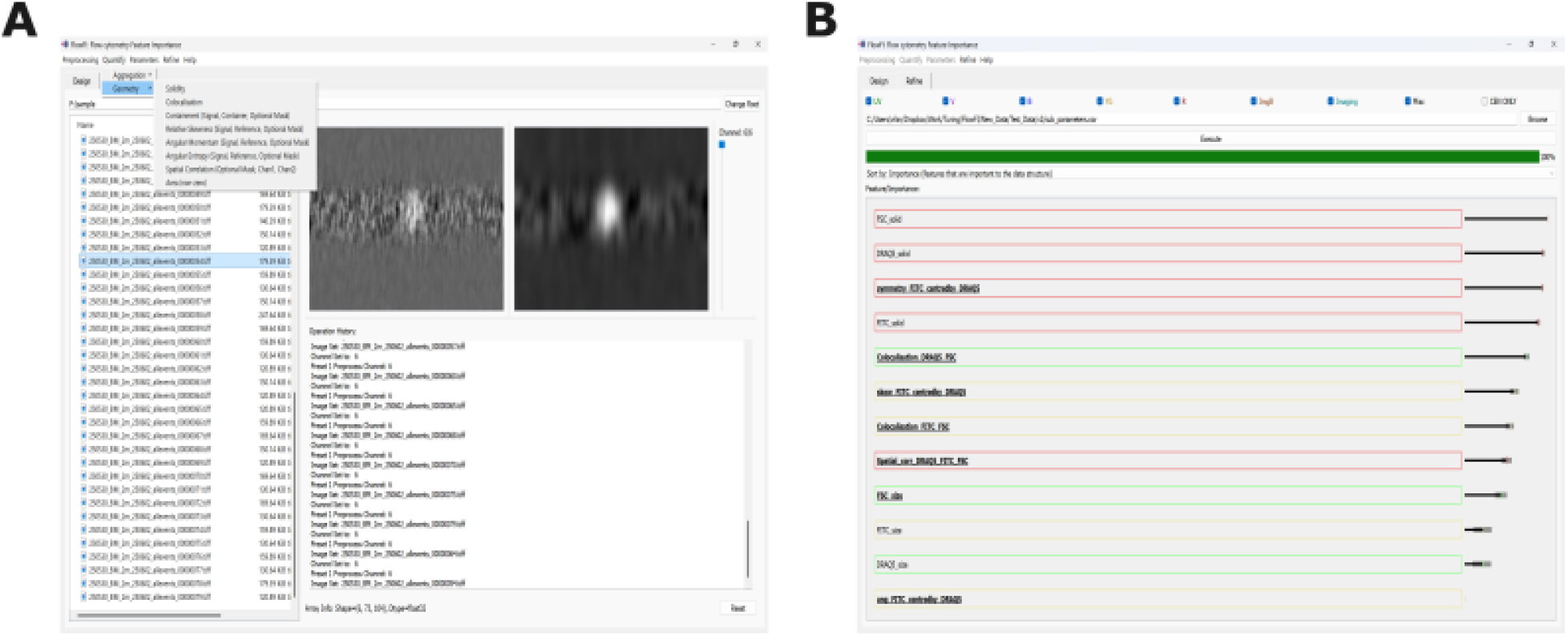
User interface for FlowFI. (**A**) The *Design* tab provides methods for image processing towards parameter quantification based on one or more image channels. Preprocessing steps can be grouped to provide a high level of customisation before selecting one of many quantification options; outputting the quantified parameter values in either FCS or CSV format. (**B**) The *Refine* tab allows for refinement of existing parameters using unsupervised machine learning techniques to suggest which parameters may be most useful in further analyses depending on your requirements.

### Image Preprocessing

Image preprocessing follows the order in which they are called and are dynamically reapplied automatically while browsing image files. Image preprocessing can be broadly separated into image *Filter* (e.g. Gaussian blur, cropping) and *Segmentation* (e.g. masking, labelling) operations (36,37). FlowFI includes preprocessing presets for application to the unique anisotropic noise profile that occurs during high throughput image acquisition in imaging flow cytometry (see Supplementary Table 1 for full range of preprocessing operations and their inputs).

### Quantification

FlowFI employs a range of methods for quantifying morphological features from cell images. These fall into two broad types, single channel quantifications involving only one image channel and multichannel quantification for features combining properties across multiple channels. At present, multichannel features are particularly underserved using available software. These are designed to work in conjunction with a preprocessing strategy, for instance quantifying the area of a given mask is likely to work best after appropriate noise reduction.

Multiple single channel quantification methods exist as part of FlowFI. Some of these morphological measures are not presently available as part of parameter calculations on any commercial software to our knowledge, including: solidity, a measure of the convexity of the objects appearing in the image, and symmetry, relative distribution of pixel intensities around a centre. See Supplementary Section 3 and Supplementary Table 2 for a full list and description of parameters, which can broadly be split into geometric (shape and relative spacing), and aggregation-based (measures based on counts of pixels or unique labels).

Multichannel quantification combines data from multiple channels into a single measurement, a feature that is unique to FlowFI as an imaging flow cytometry platform. These methods include comparative methods like spatial correlation as well as reference methods that take one image as reference (e.g. the mask defining the cell or centre of a cell). For a full list see the software documentation (or Supplementary Table 2).

### Extraction

FlowFI utilises Flowkit (38), an open source python toolkit for the manipulation of flow cytometry formats in order to both read and write flow cytometry standard FCS files, allowing for novel parameters to be appended to existing datasets for reimportation into standard flow cytometry workflows. FlowFI also allows for parameters to be saved in a single column CSV file format which can be manually reimported. FlowFI allows for optional modifications to the parameter values in order to make them interpretable in standard analysis tools which expect non-negative values.

Batch extraction of preprocessed images can be performed on entire image stacks (File) or a subset of channels which allows users to cache some of the image preprocessing steps they performed for later use as well as enable interoperability with existing image analysis software such as ImageJ.

### FlowFI: Feature Importance Workflow

Flow cytometry is an ultra-high-throughput technique that requires workflows that can scale from hundreds to millions of events and images (39). Methods need to be able to scale to these large datasets without an unsustainable increase in processing time while running on conventional laboratory work stations. In addition, the flow cytometry data is often high dimensional involving tens to hundreds of parameters. These constraints mean that data and time efficiency is required. We also seek to provide a comprehensive overview of the structure of the dataset by generating an overview of feature clusters and central features (hubs) alongside importance scores to help contextualise these scores (see Supplementary Section 2).

Importantly, the majority of feature selection methods are computationally expensive and scale poorly with sample size (40) or complexity of the data model (23). This results in methods that do not function well as part of an iterative experimental or clinical workflow. Here, we investigate a simple modification to the sampling regime in the flow cytometry feature selection pipeline which can limit computational overhead while potentially improving accuracy and robustness of ranked feature scoring (see Supplementary Section 2). In addition, we propose normalisation of all scores to improve comparability across populations or experiments.

Lastly, in order to capture the full range of what users may desire in an important feature, we provide multiple manifold learning-based measures and provide a subset of these to the user to cover a range of local and global properties of the dataset (31,35,41,42). These include using Self-Organising Maps (SOM) in combination with markerwise enrichment – notably implemented within FlowJo as FlowSOM (23) and Marker Enrichment Modelling (MEM) scoring (43)), as well as Mutual Information (MI) regression and a linear approach based on linear Principal Component Analysis (PCA) and Laplace-Scoring (LS). FlowFI currently excludes the proposed UMAP-based score (see Supplementary Section 2 for further details) due to its computational intractability, but this may be included in a future release. Fig. 3B shows how the output of the feature importance scoring is visually presented in FlowFI. Each Feature importance measure is an estimate of the relative importance of a given feature relative to others in the same data set. This Relative Importance (RI) scoring is achieved by max-min normalising the scores on a 0-1 scale for interpretability. FlowFI simultaneously provides visual feedback to the user by computing clusters of similar features based on a consensus k-medoids and Leiden modularity-based clustering procedure that also highlights the most representative features in each cluster (44–47). FlowFI also provides functionality to save and compare feature importance across datasets that share some or all of the same parameters, for direct comparison of how importance scores may vary between populations or gating strategies.

## Results

The utility of FlowFI was assessed using a dataset consisting of standard spectral and image-derived flow cytometry parameters produced using the BD FACSDiscover S8 as well as novel parameters derived directly from image data produced alongside existing parameters. The data was derived from bone marrow samples which had been specifically enriched for megakaryocytes (MKs), a rare cell type with a highly variable morphology owing to the unique level of ploidy variation within the same lineage, ranging from 2n to 128n. That means a standard cell doublet exclusion gate does not efficiently remove cell aggregates. Described size and shape variability, low cell frequency, and fragility (tendency to fragment cytoplasm that associates with other cells) all make MKs incredibly difficult to isolate from other cell types without implementing further quality control, such as with imaging.

The analysis has two parts and is designed to comprehensively test FlowFI’s functionality. After initial gating for MK enrichment, we tested whether novel parameters produced from raw image data produced on the BD FACSDiscover S8 could be used to significantly improve the refinement of the sample by removing false positive events, as defined by consensus expert labelling. This process reveals the complexity of defining a set of heuristics that fully captures the morphological heterogeneity of the MK lineage. The novel gating strategy utilising novel parameters was performed on the refined population using the original gating strategy with standard parameters. All gating was performed manually by experimenters in FlowJo, a standard software package for defining flow cytometry gating strategies.

Secondly, the robustness of FlowFI’s feature importance workflow was tested using the actual novel parameters derived from image data which were chosen for gating. A successful importance measure is one that places these parameters near the top when ranking by importance. This two-step approach tests the entire pipeline from image acquisition through novel parameter design to parameter refinement. It is meant to mimic the proposed workflow for FlowFI and its integration into the existing flow cytometry ecosystem.

### Single megakaryocyte selection

Bone marrow was isolated from 14-week-old C57Bl/6 male mouse femurs, tibias and pelvis after brief centrifugation. Bone marrow cells were filtered through a 70 μm strainer (PluriSelect) to remove debris. Red blood cells were lysed by incubation in ammonium chloride solution(Stem Cell Technologies). Megakaryocyte enrichment was performed by immunomagnetic negative selection using the EasySep Mouse Hematopoietic Progenitor Cell Isolation Kit (Stem Cell Technologies) following the manufacturer’s instructions. Megakaryocytes were stained with lineage-specific marker CD41-FITC (48) (Miltenyi) and DNA dye-DRAQ5 (49) (Thermo Scientific) and analysed with the image-based flow cytometer BD FACSDiscover S8 using BD FACSChorus v6.1 for initial gate design (BD Bioscience).

For the initial dataset 50,000 events were recorded and for the validation dataset 100,000 were recorded. Initial flow cytometry data analysis was conducted with FlowJo v10.10 software with BD CellView Lens plugin (BD Biosciences) to select for CD41 and DRAQ5 positive cells. Further gating to isolate MK from other lineage cells was performed using a combination of standard parameters available from the BD FACSDiscover S8 platform and standard imaging parameters Diffusivity (DRAQ5) and Radial Moment (CD41) were used in an additional gate to further refine the samples.

**Table 1.**
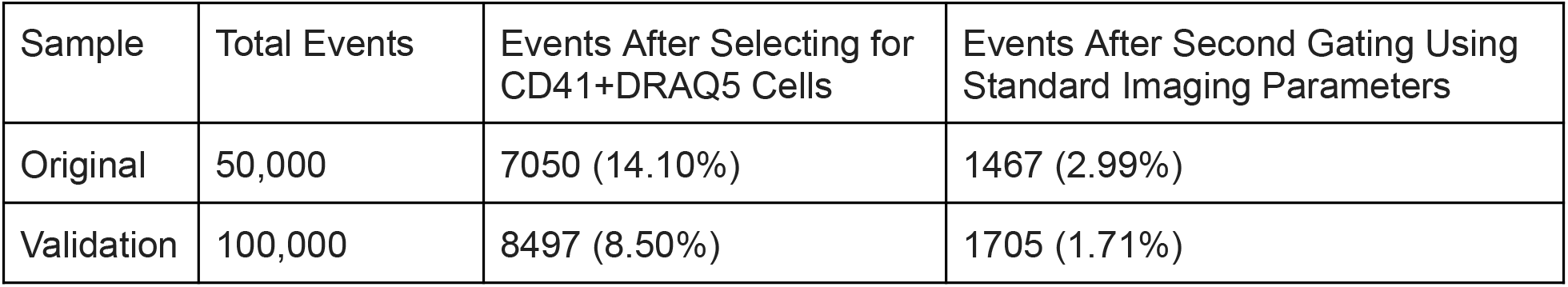
Data for the original and validation set for parameter testing and method refinement..

| Sample | Total Events | Events After Selecting for CD41+DRAQ5 Cells | Events After Second Gating Using Standard Imaging Parameters |
| --- | --- | --- | --- |
| Original | 50,000 | 7050 (14.10%) | 1467 (2.99%) |
| Validation | 100,000 | 8497 (8.50%) | 1705 (1.71%) |

MK positive events were determined by visual inspection of event images by three independent experimental raters. MK cells were further divided by ploidy using the DRAQ5-A parameter.

Inspection of each cell image stack was performed in BD CellView Lens utilising both staining (CD41 and DRAQ5) and label-free channels (see Supplementary Fig. 1). All raters were assessed based on their inter-rater agreement scores. Fig. 4A shows the agreement between reviewers (raters). The diagonal shows the mean agreement between the rater and the two others with Rater 1 having the highest mean inter rater agreement.

**Figure 4.**
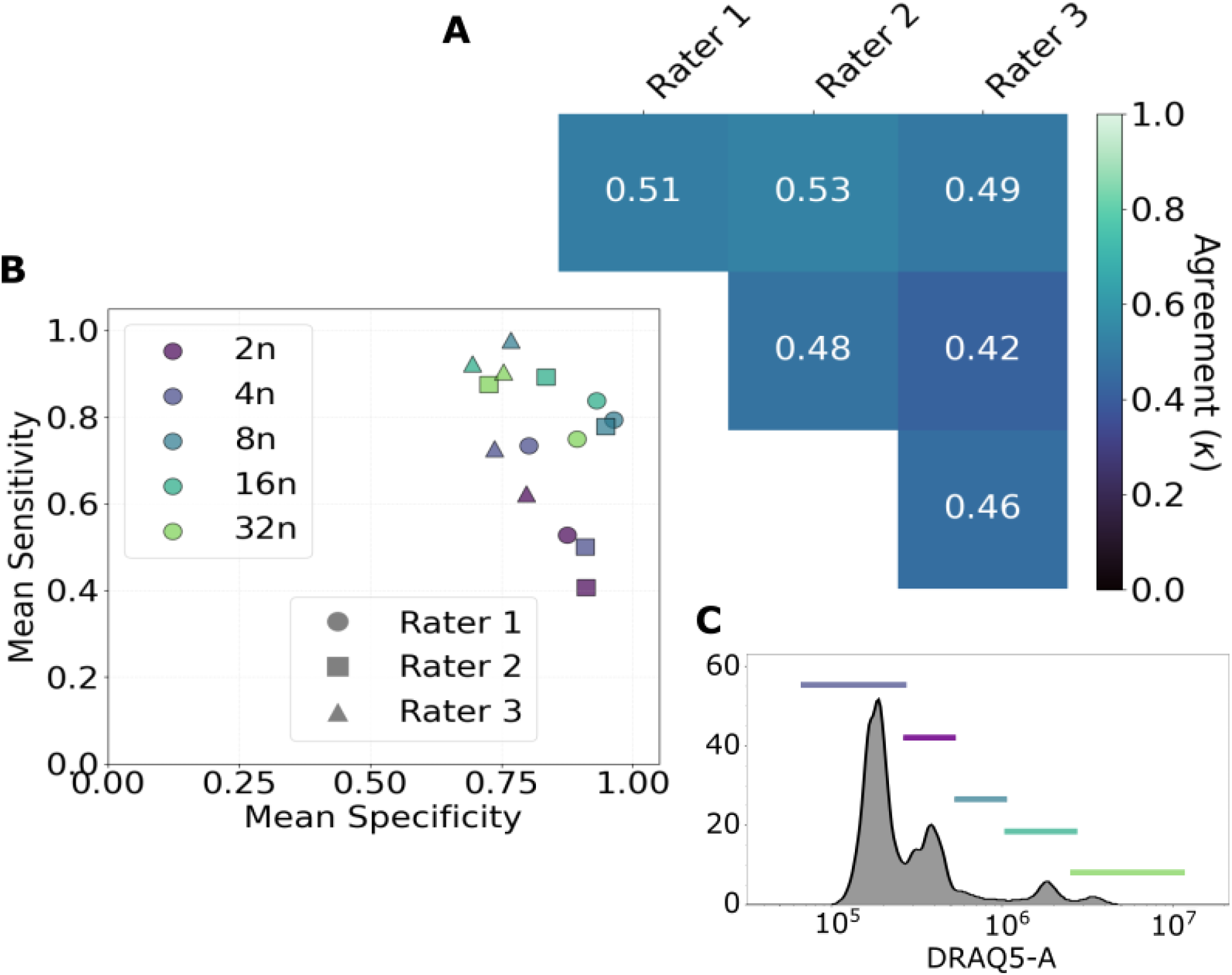
Challenges of MK classification. (**A**) Plot showing the agreement (measured using Cohen’s Kappa) between individual raters using imaging data to confirm positive MK cells (a score between 0.4 and 0.6 is generally considered moderate). The diagonal shows the mean agreement between that rater and the other two. Rater 1 shows the highest level of mean inter rater agreement. Rater 1 also has the highest consensus agreement of 0.83. (**B**) Plot of mean pairwise specificity and sensitivity. This plot shows that raters tend to agree on true negative labelling but some uncertainty exists for true positive events especially for low ploidy cells. (**C**) The histogram shows the distribution of cells by ploidy with classes defined by coloured regions across the distribution of DRAQ5-A for DNA staining area (an existing parameter). The largest number of events fall into the 2n class.

Agreement is generally low to moderate showing that the task of discriminating MK cells is complex with a high level of noise. Methods and additional parameters that can help to delineate the differences between cells are therefore desirable. Fig. 4B shows how this complexity is realised when cells are separated by ploidy. Mean sensitivity and specificity show how true the raters are to each other when separated by ploidy class. It shows that mean specificity (the tendency for raters to agree on what is not a MK tends to increase with ploidy). This is likely due to the presence of doublets and triplets which appear as high ploidy cells as well as an increase in signal strength as cells become larger, thereby indirectly contributing to an increase in the signal to noise ratio at higher ploidies. Fig. 4C shows the distribution of the DRAQ5-A parameter, a proxy for total DNA area used to determine the ploidy boundaries.

### Feature Choice

New imaging parameters were constructed using all available preprocessing and quantification options with the goal of covering the wide range of MK morphologies across all available ploidy levels. These included more standard features for size (area) as well complex geometric features such as solidity (the area of a mask relative to its convex hull, a measure of fragmentation). Both forward scatter, DNA (DRAQ5) and cell type specific cytoplasmic (membrane) staining (CD41) channels were used in parameter construction including using multichannel quantification features, notably symmetry, angular momentum, and spatial correlation which all use one or more channels or channel-derived masks to compute a summary. For a full list of novel designed parameters along with a description of each, see Supplementary Table 3.

Using the export functions (*Parameters > Export to CSV*) parameters were saved and then imported into FlowJo for manual gating. Gating was performed prior to true positive MK labelling but after gating using standard parameters, with a single additional gate chosen based on two parameters DNA solidity (DSD), the relative fragmentation of DNA, and cytoplasmic symmetry, around the DNA signal (DRAQ5 for DNA) centroid (SYCybyD). The resulting gate was recreated to fit the validation set (see Supplementary Fig. 1).

### Testing the Effect of Feature Choice on Sample Purity and Yield

Flow cytometry is a high throughput method that can efficiently enrich for rare cell types, by reducing the number of false positives (impurities). Additional enrichment can also be performed such as manual picking of true positive cells with the help of imaging data. The reduction in false positive rate (FPR) is equivalent to an increase in specificity which we tested using McNemar’s test for paired differences in false positive events. The initial FPR was high at 71.0%. Fig. 5A shows the improvement in FPR across ploidy classes upon inclusion of the novel parameter gate. Across all ploidies, the bespoke parameter gate eliminated 14.3% of existing false positives (p<0.001, McNemar), bringing some ploidy classes to majority true positive levels (notably for 8n cells and above). While the majority of removed cells were smaller 2n events, the higher proportional changes were for higher ploidies. Though the 16n class was not significant itself after Benjamini-Hochberg correction (likely due to the lower absolute count for higher ploidy cells), the higher proportional reduction for high ploidy cells is likely due to the use of symmetry and solidity parameters in the gate to distinguish multiplets, with large total nuclear area, from true high ploidy events. The overall reduction in false positives also held for the validation set (p<0.001, McNemar’s test), with the most significant reduction again for the 2n class.

**Figure 5.**
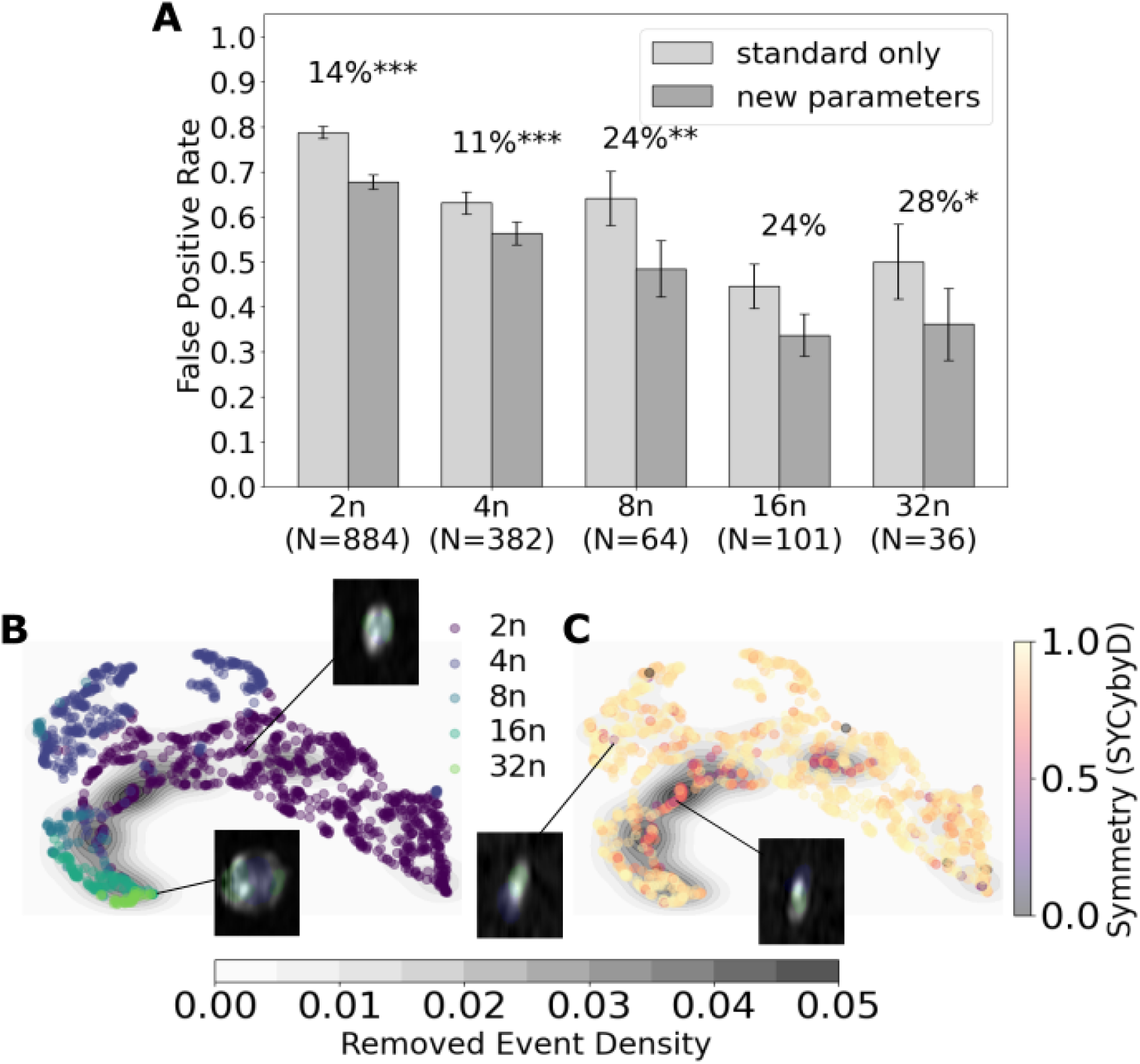
Relative improvement when gating using novel parameters. (**A**) False Positive Rate is shown when defining a gate using standard parameters (light grey) versus an additional gate using new parameters (dark grey). Change in specificity is shown across ploidy classes with the largest absolute decrease in false positives coming from the largest 2n class. Significance was calculated using McNemar’s test for paired changes in the proportion of false positives with or without the additional gate with Benjami-Hochberg correction. The overall reduction was significant at the p<0.001 significance level which was confirmed for the validation dataset (See Supplementary Fig. 3). (**B**) This UMAP based on the original standard gating parameters shows events separated by ploidy in parameter space with examples for high and low ploidy. The majority of events removed by the new gate (grey density plot) appear to be at the boundary between low (≤8n) and high (≥16n) ploidy cells, suggesting the new gate primarily removes ambiguous events with intermediate morphology. (**C**) The gating parameter SYCybyD measures the symmetry of cytoplasmic signal around the nucleus. It suggests that SYCybyD is predictably lower at the class boundary due to the presence of irregular doublets, triplets and nuclear fragments around the nuclei (two low symmetry examples are shown).

Imaging flow cytometry using the BD FACSDiscover S8 is a high throughput method that makes acquisition of large samples relatively quick and low cost even when cell types are rare or gating is strict. However, significant decreases in the yield of true positive cells would significantly reduce the statistical power of downstream analyses or force experimenters to spend time acquiring larger samples or running costly additional experiments. We tested whether there was a significant cost in the loss of true positives (decreased sensitivity) at a predetermined 2% threshold using McNemar’s test. We found no significant evidence supporting a greater than 2% decrease in sensitivity for any ploidy class (after Benjamini-Hochberg multiple testing correction). A slight decrease in sensitivity was observed for the validation set of no more than 6% across any one ploidy class. This suggests that given the significant increases in purity (specificity) the decrease in overall cell yield is likely worth it in this specific refinement for the majority of use cases.

The UMAP in Fig. 5B shows that ploidy classes clearly share similarities across the standard gating parameters used for the embedding with examples shown of a high and a low ploidy cell (including nuclear and cytoplasmic mask). The images exemplify the overall cell size differences between classes. Areas of highest removal density (grey) show that the majority of removed/excluded cells occur at the high-low ploidy boundaries. This supports the multiplet contamination hypothesis since multiplets likely have spectral and visual characteristics that are in some sense intermediate between high and low ploidy cells. Fig. 5C shows the novel symmetry (SYCybyD) parameter used in the novel parameter gating strategy. The low symmetry events also cluster around the ploidy class boundary.

### Feature Importance Benchmarking

The previous section established the utility of novel parameters in reducing sample impurities for MK purification. We now show how this parameter search may have been sped up using the feature importance methods available in FlowFI. In total 5 methods were tested that represent a range of approaches to unsupervised feature selection. Tab. 2 shows the 5 methods that were benchmarked which cover a range of approaches from local to global structure preservation to information theoretic, linear, non-linear and topological measures of importance. FlowFI’s default feature importance measure utilises Laplace-Scoring, which measures the degree to which a feature is in alignment with the local neighbourhood structure of the nearest neighbour graph or manifold based on the data (31). One of these is the novel, UMAP-KLD measure (uRI) which is meant to capture the parameter intuitions commonly derived from such low dimensional representations of flow cytometry data in an experiment (see Supplementary Section 2.2 for importance score definitions).

**Table 2.** Feature importance methods split by the focus of the model (global vs local structure preserving features) and definition of importance used.

| Method | Local-Global Structure | Importance Methodology |
| --- | --- | --- |
| Laplace-Scoring (lsRI) (31) | Intermediate | Neighbourhood preservation |
| PCA (pRI) (42) | Global | Linear variability explained |
| UMAP-KLD (uRI) (50) | Global | Contribution to manifold compression |
| Cluster-SOM (fRI) (41) | Local | Cluster identity preservation |
| Mutual Information (miRI) (35) | Global | Maximum informational dependence |

All methods were compared using two different sampling approaches, fitting the model to the full dataset or repeated fitting of the model using a smaller sub-sample and aggregating the results over multiple (B=1000) bootstraps. Fig. 6A shows Mean Average Precision (MAP) scores, a rank reliability measure of selection accuracy (see Supplementary Section 2.3). MAP scores were uniformly higher when bootstrapping was performed. The highest MAP scores are for bootstrap Laplace Scoring, 0.833 and bootstrap PCA-based RI, 0.700 which are both non-local, non-cluster focused methods. The novel UMAP-KLD method performed moderately at 0.583 for bootstrapping with all other methods performing below the general cut-off for moderate agreement < 0.4. Fig. 6B shows the computational cost of RI acquisition summed over all samples and features. The full dataset calculation tends to suffer from the poor scaling properties of most feature selection methods (e.g. O(n^2^) for UMAP, MI and LS which use the k-NN graph algorithm). Fig. 6C shows a range of RI values for all methods (using bootstrap aggregation). The majority of methods agree on a high RI for SYCybyD with disagreement for the relative importance of DSD.

**Figure 6.**
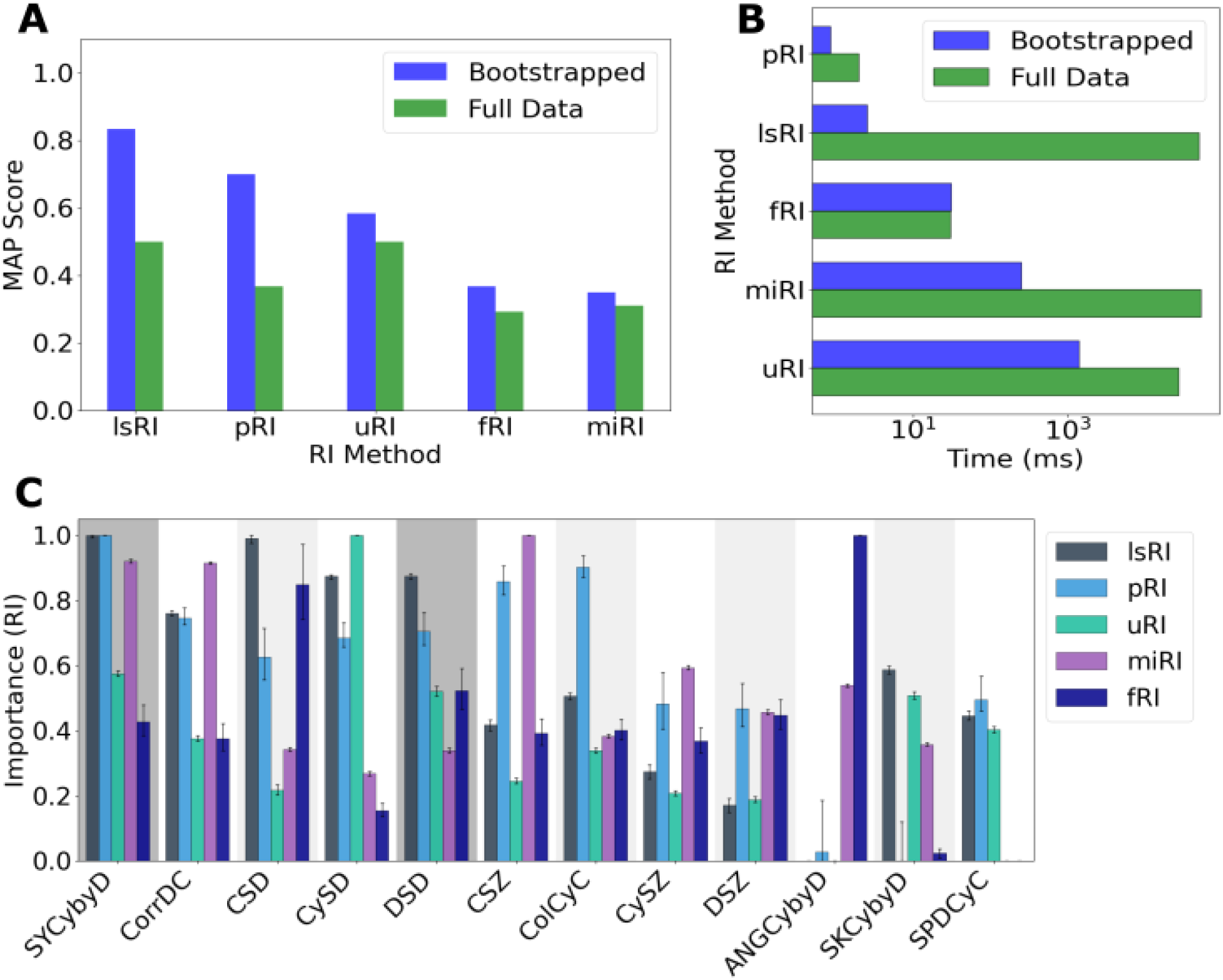
Testing of unsupervised feature importance measures using experimentally confirmed parameters. (**A**) Mean Average Precision (MAP) scores are shown across the measures when using subsampled bootstrapping versus the entire dataset. (**B**) Log-scale bar plot of compute time taken to calculate each RI measure using either bootstrapping (B=1000) or the full dataset. Since many of these methods rely on algorithms (such as k-NN graph construction) with a O(n^2^) time complexity over the sample size, n. (**C**) Shows the performance using a range of unsupervised learning measures of Relative Importance (RI) of the novel parameters. Dark grey blocks show which novel parameters were selected by experimentalists in the gating experiment. High RI indicates a high confidence in this feature across bootstrap subsamples with error bars indicating the 95% empirical CI. SYCybyD was measured to have a high importance across most measures while DSD is high only for lsRI and pRI measures.

## Discussion

FlowFI provides a fully open source and interactive platform for the emerging field of IFC and flow cytometry feature analysis more broadly, addressing gaps that complement existing workflows and are designed to be compatible with available flow cytometry software for both sorting and analysis, focusing on parameter design in emerging areas such as intracellular and intercellular interaction analysis (51), and label-free sorting (52). FlowFI provides an image preprocessing and multichannel parameter quantification platform for novel parameter construction and full exploration of image space usable by both wet lab and computational specialists. FlowFI focuses on providing bespoke preprocessing options tailored to the specific image artefacts common in high throughput image acquisition with IFC (53), enabling both better feature design and improved signal to noise ratio for downstream image analysis or machine learning-based models (54). FlowFI combines these image analysis features with ensemble-based manifold learning for novel parameter comparison and refinement of the parameters produced through exploration.

MKs represent an immunologically (55), and haemostatically important, highly heterogeneous (56,57), and rare population of bone marrow cells. Recent reports have shown coexistence of numerous subpopulations that undergo considerable morphological changes across multiple axes, notably polyploidisation but also differentiation and functional specialisation within specific ploidies (18,57). This makes these cells a unique challenge to sort as a single population using existing parameters. These factors as well as other idiosyncratic characteristics such as the tendency to form cellular aggregates (multiplets) with other haematopoietic cells (58), and emperipolesis, the temporary engulfment of neutrophils by MK cells (59,60), contribute to the limited characterisation of the MK lineage, both morphologically and transcriptionally (19,61,62). We used FlowFI to eliminate false positive events with low symmetry and high nuclear fragmentation (low solidity) from an MK enriched BM sample. Eliminating significantly more impurities than would have been possible with standard imaging and spectral parameters, resulting in improved purity across subpopulations at a relatively low cost to specificity or yield. Though the majority of removed events were classified as 2n cells, the highest overall improvements were in the higher ploidy classes. This difference can be partially explained by the lower signal to noise ratio for smaller, low ploidy events and the higher prevalence of multiplets, and other fragments removed by the new gate.

We tested FlowFI’s feature importance and parameter refinement tool across a range of standard measures, including those based on measures in existing platforms such as FlowJo, using analogues to existing methods such as SOM and PCA (23,42). In addition we tested a measure based on the novel application of the UMAP algorithm (50) to capture the importance placed on clustering within embeddings in the interpretation of cell populations. FlowFI’s ensemble bootstrap subsampling approach showed improvements in both performance (as measured by Mean Average Precision) and computational efficiency over what can be achieved one learning directly from the full dataset, improving scalability of these methods to high throughput data. One possible explanation for this is that bootstrap aggregation serves as a regulariser, ensuring that features must be stably predictive of the manifold substructure in order to score high (63,64). This serves to mitigate the influence of noise that may dominate the manifold structure when inferring from the full dataset. The apparent success of methods like Laplace Scoring and PCA, and underperformance of MI and established methods like SOM in unsupervised feature selection may be due to those methods fairing better for higher dimensional data. It could also be due to the density dependence of the latter two methods (35,65), making them more sensitive to the relatively dissimilar false positive cells, which are composed of anything from small cell fragments to large heterogeneous cell aggregates.

FlowFI’s feature importance pipeline circumvents much of the computational cost of processing a large high dimensional flow cytometry dataset all at once by sub-sampling the data repeatedly. This can lead to potential blindspots when the subpopulations of interest are very small in size (low signal-to-noise) as the subsampling can serve to smooth out the underlying signal. Arguably, this is a problem for unsupervised methods more generally (66), however FlowFI allows for bootstrap parameters to be changed to account for differences in composition. FlowFI also offers an adjustable early stopping threshold (see Supplementary Section 2.5) to be used if computational cost or over smoothing may be a concern as well as a coverage estimate based on subsample-to-sample size ratio (67,68). In future, we plan to add additional group feature selection measures that more closely match the goals of parameter-based gating (69), and make further improvements through hardware specific optimisation of both image and feature analysis pipelines (70). FlowFI provides a useful testing ground for future developments in this space with planned improvements to its feature and manifold learning capabilities, as well as improved integration with common research image analysis platforms like ImageJ (71).

Megakaryocytes are highly heterogeneous and their large size provides a unique advantage for morphological profiling and testing of FlowFI as a platform for diverse morphological parameter design. The scale of IFC enables identification of rare and infrequent subtypes as well as tracking of population-level shifts in morphological traits as a function of disease or other stress conditions (inflammation, aging, dietary intervention). In particular, single cell transcriptomics can help us interpret the meanings of these morphological population-level shifts and create a two-way mapping from morphology to expression. Crucially, though image-based parameters and visual inspection are prone to error due to the prevalence of high throughput image acquisition artefacts, these steps act as an invaluable quality control step for future work in morphotranscriptomics and transcriptomics as a whole.

## Conclusion

We introduce FlowFI, a tool for designing novel parameters specific to the experimentalist or clinicians needs and addresses the growing gap between standard morphological and spectral features and the range of possible morphologies and configurations studied across the field. FlowFI also offers a less costly alternative method for feature selection based on a computationally efficient sampling approach resembling an average of experts framework, which provides scalable insights into feature identity and utility within a range of possible interpretations. Beyond megakaryocyte biology, this approach has broader potential as a label-free, cell type-agnostic strategy for profiling diverse biological kingdoms and in providing new axes along which to explore pathological cell presentation for diagnosis. In particular, it may be especially useful in under-characterised or underserved systems such as protists, pollen, and plant protoplasts, where the availability of specific antibodies and thus the capacity for marker-based enrichment, is limited. Here, morphology or autofluorescence-based sorting offers a practical alternative that could greatly expand imaging flow cytometry’s applicability to complex microbial, environmental, and biomedical contexts.

## List of Abbreviations

IFC: Imaging Flow Cytometry
IACS: Image Assisted Cell Sorting
OFDM: Orthogonal Frequency Domain Multiplexing
MK: Megakaryocyte
BM: Bone Marrow
DSD: DNA (DRAQ5) Solidity
SYCybyD: Symmetry of Cytoplasm (CD41-FITC) around the DNA (DRAQ5) centroid
UMAP: Uniform Manifold Approximation and Projection
KLD: Kullback-Liebler Divergence
PCA: Principle Component Analysis
SOM: Self Organising Map
MEM: Marker Enrichment Modelling
RI: Relative Importance
FlowFI: Flow cytometry Feature Importance
Xn: Number of complete set of chromosomes, X, found in a cell or organism (i.e. ploidy)

## Declarations

### Ethics approval and consent to participate

All animal experiments were conducted in full accordance with the Animals (Scientific Procedures) Act 1986 under UK Home Office (HMO) approval.

### Consent for publication

Not applicable

### Availability of data and materials

Software is available as source code on a MIT CC BY license on github and also includes binary standalone installable versions of the software for Windows along with comprehensive documentation. Documentation is also available on execution from the Help menu.

