## Supplementary Information for "FlowFI: an interactive graphical software package for bespoke design of imaging parameters in flow cytometry to explore morphological diversity in bone marrow megakaryocytes"

This supplementary information document covers the implementation, mathematical and validation details.

### 1 Preprocessing Options

| **Name** | **Description** | **Parameters** |
|---|---|---|
| Gaussian Filter | Applies Gaussian smoothing to reduce noise | $\sigma$ |
| Crop | Removes pixels from image borders | Edges (px) |
| Rescale | Resizes the image dimensions | Scale (X,Y) |
| Mask Otsu | Binary thresholding with morphological cleanup | None |
| Label Image | Labels connected components in a binary image | None |
| Segment | Separates touching objects using watershed | None |
| OMDF Preset | Composite image preprocessing preset | None |

Table 1: Summary of available preprocessing operations in FlowFI.

Table 1 shows the available image preprocessing options in FlowFI. Implementation information is documented in Python's scikit-image for Gaussian filtering, Otsu thresholding, and watershed segmentation [1].

#### 1.1 Preset for Waveform-based Image De-noising

The the waveform-based image de-noising preset option is specifically designed to deal with the noise signatures present in the Orthogonal Frequency Domain Multiplexing (OFDM) approach used by the BD S8 and A8. OFDM for image reconstruction involves splitting of a single laser beam into many beamlets. The beamlets are arranged in parallel spatially and operate at different frequencies, designed in such a way as to be mathematically orthogonal in frequency space, which means they can be clearly separated on the detector side. This allows for spatial reconstruction of the image [12]. The single origin plane of the beamlets results in specific anisotropic noise signatures which in FlowFI are handled using a series of scaling and filtering steps for computational efficiency. In FlowFI the *OMDF Preset* is composed of a series of preprocessing steps:

- $LEFTCROP(10)$ - To remove reconstruction artifacts
- $RESCALE(X * 1, Y * 0.25)$ - Rescale to approximate original pixel sizes for BD S8
- $GAUSS(\sigma_x = 5/2, \sigma_y = 4/5, K = (5, 3))$ - Anisotropic noise reduction
- $RESCALE(X,Y/PAR^*)$ - Equalize aspect ratio
- $GAUSS(\sigma = (3/2, 3/2), K = (3, 3))$ - Isotropic noise reduction

*$PAR$ is the Pixel Aspect Ratio specific to the cytometer set-up (a standard value on BD S8 machines is 1.2).

### 2 Importance Measure Evaluation

Data, $X$, is standard scaled with zero mean unit variance. Bootstrapping is performed over all $n = 200$ for all methods over $B = 1000$ runs, providing the RI measure which is a mean over all $B$ samples so that

$$\bar{I}(m) = \frac{1}{B} \sum_{b \leq B} I(m; b)$$

where $I(m; b)$ is the importance score measured for subsample $b$ of size $n$. The RI is then

$$RI(m) = \left( \frac{\bar{I}(m) - \min(\bar{I})}{\max(\bar{I}) - \min(\bar{I})} \right) \tag{1}$$

#### 2.1 Empirical Confidence Interval Calculations

Confidence intervals were calculated by sampling importance vectors $I(\cdot, b)$ from the $1 \leq b \leq B$ repetitions. Mean values were recalculated with replacement $K = 10,000$ times resulting in an empirical distribution of bootstrap $\bar{I}$ estimates, $\tilde{I}$. The relative importance bootstrap $\tilde{RI}(m)$, were then used in determining the 95% CI for the observed $RI(m)$. Importantly, since samples were based on $I$, which can have support in all of $\mathbb{R}$, the empirical $RI$ estimates were clamped at the top and bottom using the original maximum and minimum estimates to ensure consistency of the estimator

$$\tilde{RI}(m) = \text{clamp}\left( \left[ \frac{\tilde{I}(m) - \min(\bar{I})}{\max(\bar{I}) - \min(\bar{I})} \right], 0, 1 \right). \tag{2}$$

#### 2.2 Importance Measures

We briefly touch on the different importance measures used in feature selection

##### 2.2.1 Laplace Scoring Importance (lsRI)

Laplace Scoring (LS) importance measures how much a feature is able to preserve the local geometric structure of a data manifold. Laplace Scoring uses the $k$-Nearest Neighbour ($k$-NN) algorithm to construct a manifold reprsented by as a graph $G$ [3], here based on the dot product of feature vectors. It uses the degree matrix of $G$, $D$, to construct a normalised feature vector for each feature $m$,

$$\tilde{f}(m) = X_{\cdot m} - \left( \frac{X_{\cdot m}^T D \mathbf{1}}{\mathbf{1}^T D \mathbf{1}} \right) \mathbf{1}.$$

The graph Laplacian, $L$, is then used to measure the smoothness or agreement $\tilde{f}(m)$ between feature $m$ and the graph manifold,

$$I_{ls}(m) = -\frac{\tilde{f}(m)^T L \tilde{f}(m)}{\tilde{f}(m)^T D \tilde{f}(m)}.$$

The number of neighbours $k = 15$ is set to be consistent across all methods we apply that utilise the $k$-NN approach to manifold learning. The minus sign in front of the above ratio means that the measure is high when variability is low (high smoothness) in the feature values of $m$ when we consider the change in $m$ between neighbours in $G$.

#### 2.2.2 PCA-based Maximal Variation Importance (pRI)

Principal Component Analysis (PCA) is a common method used in dimensionality reduction and feature analysis which is available in packages like FlowJo to determine the axes of maximal variation in a multidimensional feature space. We define a measure of variability attributed to a feature $m$ using the feature loadings, $V$, derived from SVD of $X$, a standard linear measure of global variability due to $m$ [11]. The method sums all absolute loading values of $m$ up to a threshold proportion of the total variance explained, $\theta = 0.8$ (a standard value for dimensionality reduction). The Explained Variance Ratio (EVR) can be derived directly from the eigenvalues $\Lambda$,

$$\text{EVR}_m = \frac{\Lambda_m}{M}$$

and so the importance score is

$$I_p(m) = \sum_{\text{EVR}_i \leq \theta} |V_{i,m}|.$$

#### 2.2.3 UMAP Importance (uRI)

This importance metric measures the unique topological contribution to the structure of the data manifold as defined by the UMAP data matrix [9].

$$\text{BCE} = -\sum_{i,j} (P_{ij} \log(Q_{ij} + \epsilon) + (1 - P_{ij}) \log(1 - Q_{ij} + \epsilon))$$

where

$$Q_{ij} = \frac{1}{1 + a d_{ij}^{2b}},$$

and $\epsilon = 1e - 10$. and $P_{ij}$ is a (fuzzy) nearest-neighbour adjacency matrix with standard parameter $k = 15$ [cite]. The importance score is then defined the difference in Binary Cross Entropy (BCE) loss

$$I_u(m) = \Delta BCE = \text{BCE}^*(m) - \text{BCE}_{\text{base}}$$

where $\text{BCE}^*(m)$ is defined as the BCE value obtained when the $i^{th}$ column (parameter) of $X$ is randomly permuted. This permutation has an effect on the nearest neighbour graph of $\text{BCE}^*(m)$, $P^*(m)$, so that $P^*(m)$ now includes false neighbours induced by destroying the structure feature column $X_{\cdot m}$. The measure therefore measures the change in UMAP compression efficiency (goodness of fit) when structure, specifically data structure owing to feature $i$ is lost.

#### 2.2.4 Mean Mutual Information Importance (miRI)

Mutual information regression uses the k-NN algorithm to assess the amount of information gained about a univariate variable $y$ by observing another univariate variable $x$, it is defined as

$$I(x; y) = \psi(k) + \psi(n) - \frac{1}{n} \sum_{i=1}^{n} [\psi(n(i) + 1) + \psi(n_y(i) + 1)],$$

where $\psi$ is the digamma function, $k = 15$ is the number of neighbours, $n$ is the sample size, and $n_x(i)$ are the marginal neighbour sets. These are defined with respect to the $L_\infty$ norm, the $i^{th}$

data point, and $k$ [6].

The mutual information-based feature importance is then defined as

$$I_i(m) = \frac{1}{M-1} \sum_{m \neq l} I(X_m; X_l),$$

which measures the informational dependence (or redundancy) of feature $m$ with respect to the other features.

##### 2.2.5 Self Organising Map Cluster Importance (fRI)

This measure is inspired by feature analysis tools already available in packages like FlowJo (FlowSOM). The Self-Organising Map (SOM) measure uses the SOM embedding algorithm which utilises a neural network arranged in a grid with a simple updating rule [5]. We use SOM to first determine an embedding appropriate to subsampling with $n = 200$ (and use the same for $n = N$, the full dataset), with square grid dimension $s = 5$ (a $5 \times 5$ grid shape). We use the implementation in the Minisom library [14].

In order to determine the cluster-preserving importance of feature $m$ for each subsample we assume that data is first clustered into $C = 5$ metaclusters with membership determined by agglomerative Ward clustering [16]. Membership preservation is then evaluated by the Marker Enrichment Modelling (MEM) score [2], as available in FlowJo

$$\text{MEM}(m, c) = |\text{median}(c) - \text{median}_{ref}| + \left( \frac{IQR_{ref}}{IQR_c} - 1 \right).$$

This score compares the median of feature $m$ for cluster $c$ to a reference median (derived from the full subsample). The total importance is then the sum of the MEM score over all available clusters,

$$I_f(m) = \sum_{c=1}^{C} \text{MEM}(m, c).$$

This measures total enrichment of a feature $m$ over all clusters.

#### 2.3 Measure Performance Evaluation

Mean Average Precision measures whether a given measure places the subset of relevant features near the top of the ranked measurements,

$$\text{MAP} = \frac{1}{R} \sum_{m=1}^{M} (P(m)\text{rel}(m))$$

where $R$ is the total number of relevant features, rel$(m)$ is the indicator which is one if $m$ is a relevant feature and otherwise zero, and $P(m)$ is the precision at cut-off $m$ in the ranked list.

#### 2.4 Consensus Feature Clustering and Feature Centrality

FlowFI employs a consensus clustering approach to identify groups of co-varying features as well central features in the data manifold. Consenus clustrting is an established method for mitigating the variability inherent in stochastic clustering algorithms and subsampling [7, 17].

In each bootstrap iteration, features are partitioned using the Partitioning Around Medoids

(PAM), a type of k-Medoids algorithm [10]. The k-Medoid method was chosen for its robustness to outliers compared to k-Means, as it uses actual data points (features) as cluster centres, these can be used to additionally define a consensus centrality measure over a given run,

$$\bar{H}(m) = \sum_{m \leq M} \sum_{c \leq C} \frac{\delta_c(m)}{M},$$

where $\delta_c(m)$ is the indicator function which is 1 if $m$ is the centre of cluster $c$ and 0 otherwise. When averaged over all $B$ bootstraps, the consensus centrality $\tilde{H}(m)$ measures the tendency of a feature to appear as a cluster centre, a potential supporting piece of evidence in feature selection.

For the final feature clustering, a co-occurrence matrix is constructed by aggregating the bootstrapped clusterings. Each entry $D_{ij}$ represents the frequency with which feature $i$ and feature $j$ were assigned to the same cluster across all bootstrap iterations

$$D_{i,j} = \begin{cases} 0 & \text{if } i = j \\ \sum_{b \leq B} \delta_b(i,j) & \text{if } i \neq j \end{cases},$$

where $\delta_b(i,j) = 1$ if features $i$ and $j$ are assigned to the same medoids cluster in bootstrap $b$, and 0 otherwise. This co-occurrence matrix is treated as a weighted adjacency matrix of a graph, and the Leiden algorithm is applied to this graph to maximize modularity, producing the final feature clustering. The Leiden algorithm is preference for connected communities that can occur, for instance when using the Louvain algorithm for clustering [13].

#### 2.5 Convergence Checking and Stability Analysis

To ensure the reproducibility of results without performing an excessive number of bootstrap iterations (reducing computational overhead), FlowFI provides optional convergence checking of the RI ranks based on split-half validation [8].

After a minimum number of iterations, the accumulated bootstrap results are periodically split randomly into two halves. The stability of the feature ranking and clustering is assessed by comparing the results derived from these two independent subsets.

The mean feature importance scores are calculated for both halves. The similarity between the two resulting rankings is quantified using Kendall's Rank Correlation Coefficient ($\tau$) [4]. Convergence is determined if the disagreement (defined as $1-\tau$) is below a user-defined threshold ($\epsilon$), indicating that performing more iterations is unlikely to change the ranking of features.

If the rankings have converged, the stability of the feature clusters is assessed. A consensus clustering is derived for each half (see Feature Clustering below). The agreement between the two partitions is measured using the Adjusted Mutual Information (AMI) score. [15].

### 3 Quantification Options

FlowFI uses a variety of parameter quantification options that are designed to work in combination with the existing image preprocessing steps. Table 2 shows the range of all available quantification options available in FlowFI, giving the specific number of input channels involved (in the case of multichannel quantification), as well as the numerical range of the parameter produced.

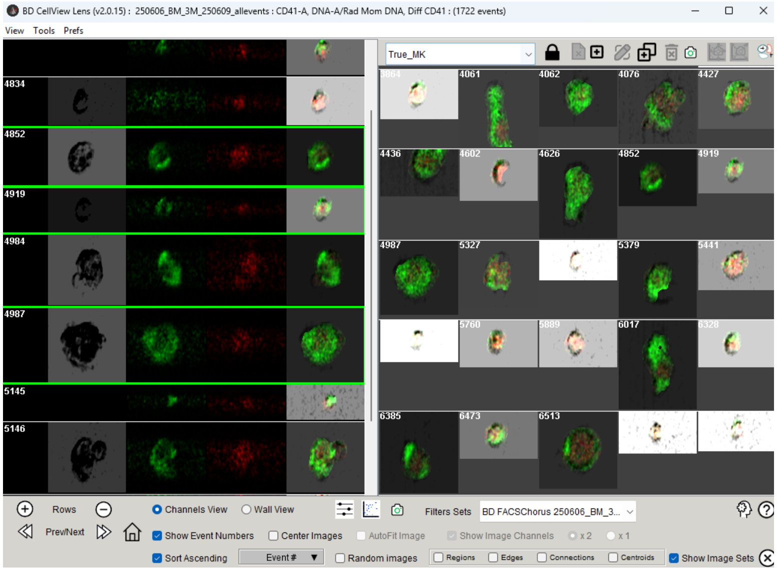


(a) Selection View

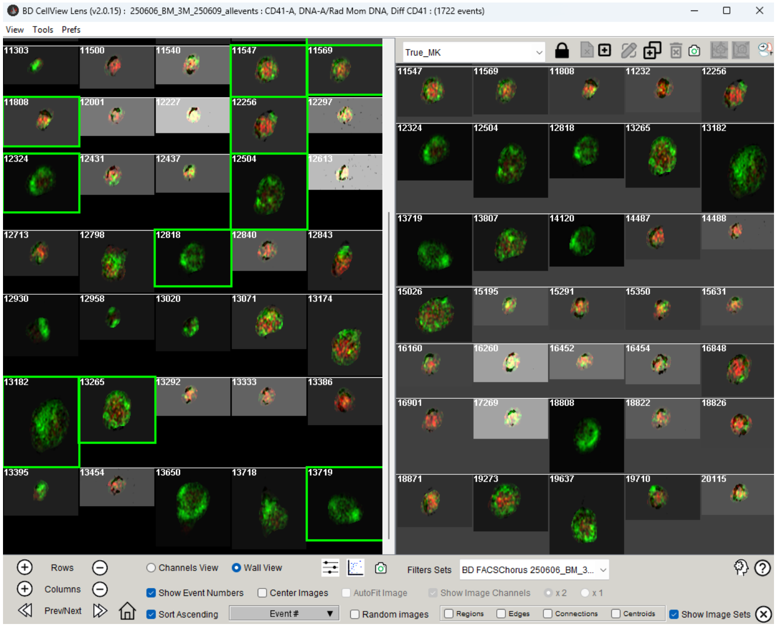


(b) Population View

Figure 1: Inspection and selection of cells in BD CellView Lens. (**A**) This screenshot shows how data from multiple channels is integrated into manual MK cell labelling. (**B**) This figure shows the full population (left) of the standard gate with selected

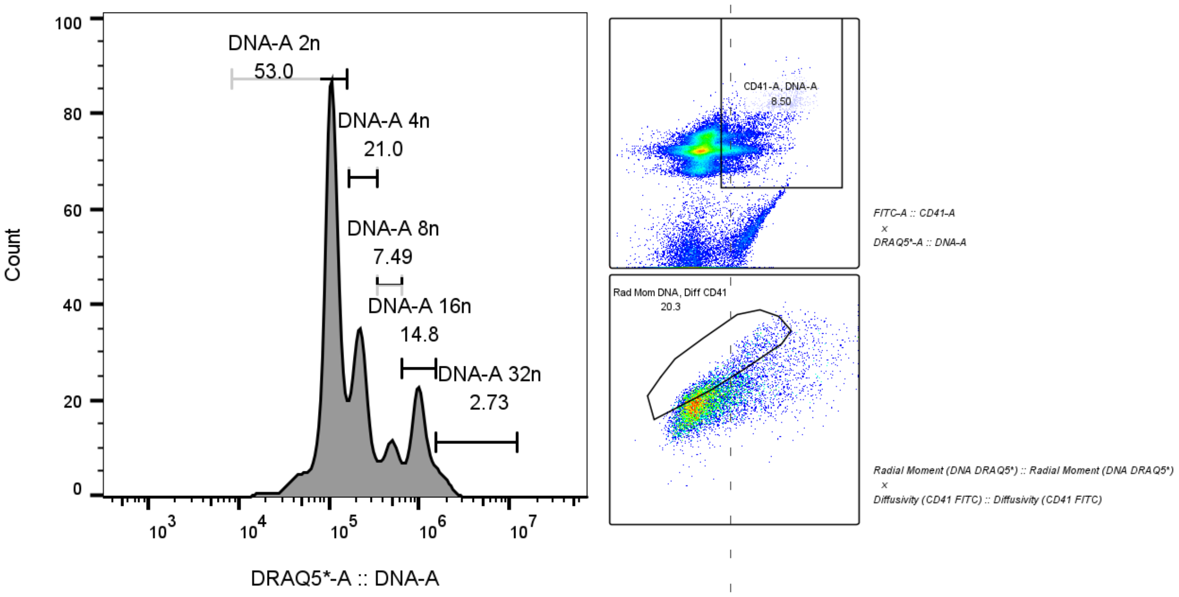


Figure 2: Initial gating strategy used in MK isolate (top and bottom right) and definition of ploidy classes (left) produced in FlowJo v10.10.

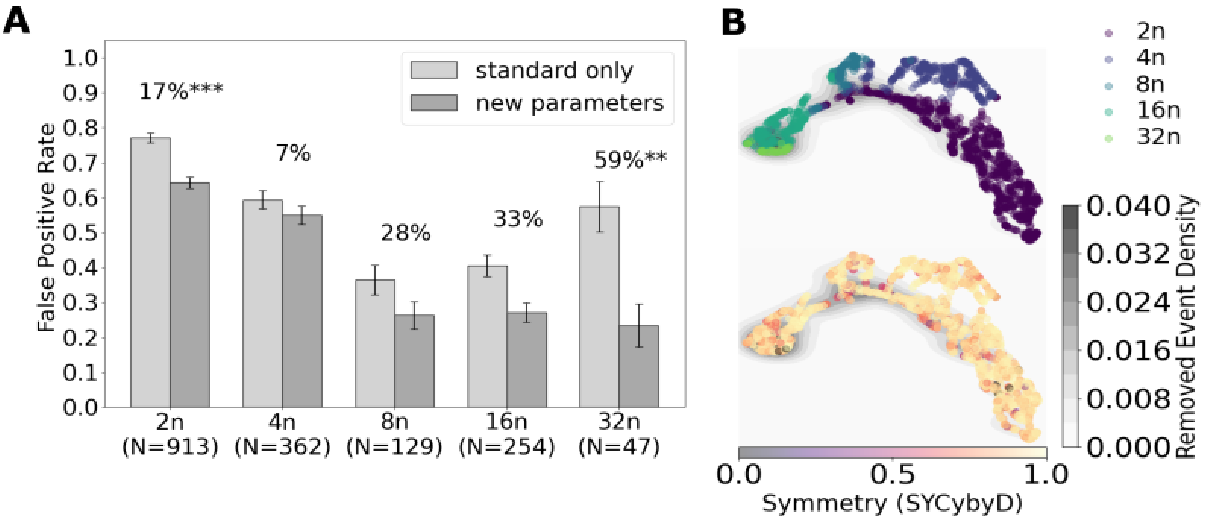


Figure 3: Validation dataset results for MK purification and false positive removal. (A) Shows the rate of false positives before and after the new parameter gate is added, again showing a significant reduction primarily in lower ploidy classes, though not all classes are significant the largest class of cells, 32n, saw a 59% reduction. (B) This figure shows a UMAP embedding of the events in the validation dataset using the original parameters, showing a similar topology and organisation by ploidy class.

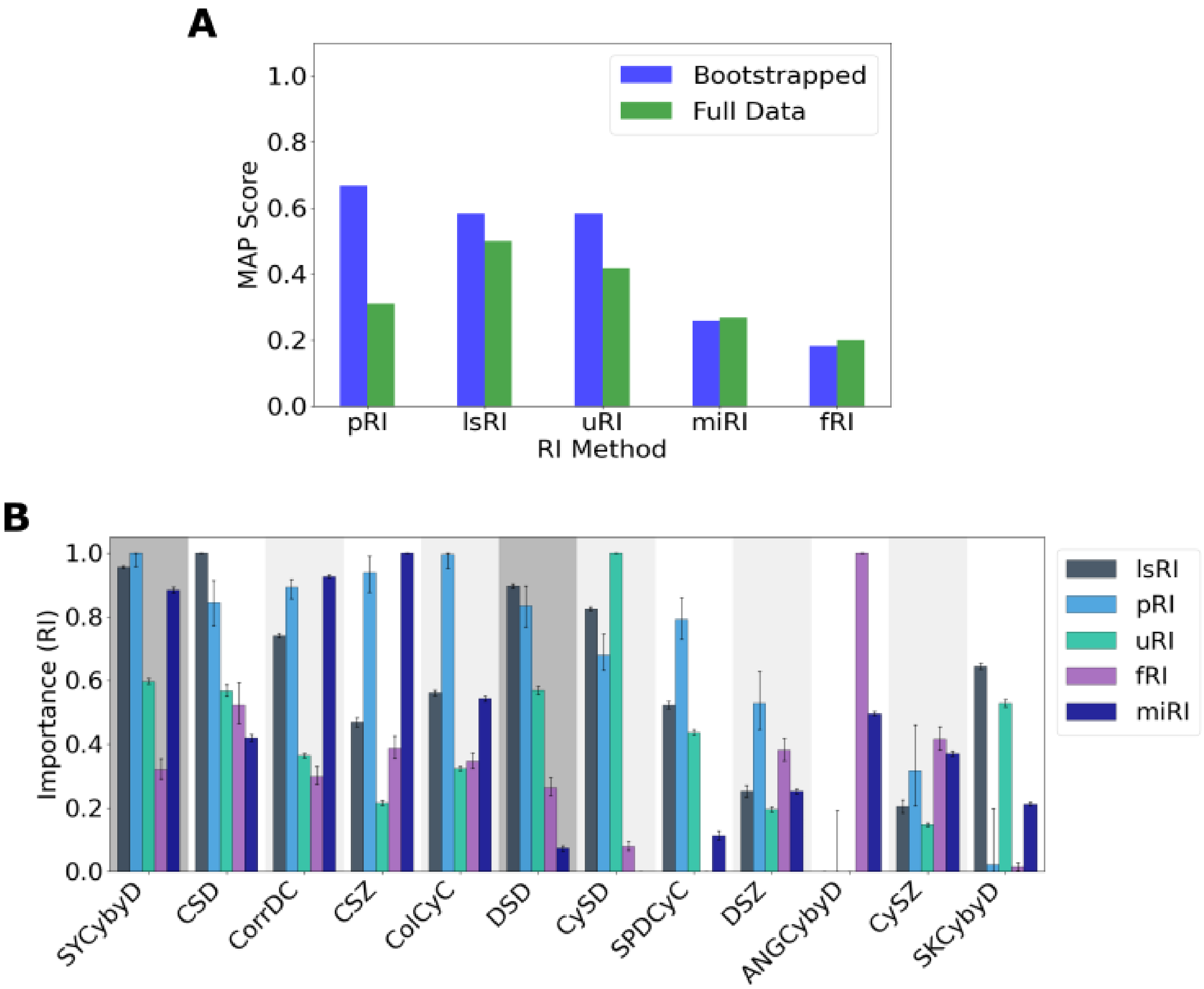


Figure 4: Validation dataset results for the refinment pipeline. (A) This figure shows Mean Average Precision (MAP) scores for each method comparing the bootstrap ensemble approach to full dataset-based feature selection. In most methods ensemble bootstrapping outperforms or is comparable to each of importance scoring methods. The linear PCA based method marginally outperforms Laplace Scoring and the UMAP method but MI and SOM methods still perform poorly for this dataset.

| Name | Description | Inputs | Range |
| --- | --- | --- | --- |
| Count | Number of unique non-zero regions | 1 | $\mathbb{Z}_{\geq 0}$ |
| Mean | Average intensity of non-zero pixels | 1 | $\mathbb{R}_{\geq 0}$ |
| Area | Total number of non-zero pixels | 1 | $\mathbb{Z}_{\geq 0}$ |
| Solidity | Ratio of object area to convex hull | 1 | $[0, 1]$ |
| Colocalisation | Fraction of signal intensity in mask | 2 | $[0, 1]$ |
| Containment | Fraction of signal inside container | 2-3 | $[0, 1]$ |
| Relative Skewness | Radial skewness from reference | 2-3 | $\mathbb{R}$ |
| Angular Momentum | Magnitude of angular asymmetry | 2-3 | $\mathbb{R}_{\geq 0}$ |
| Symmetry | Uniformity of angular distribution | 2-3 | $[0, 1]$ |
| Spatial Correlation | Pearson correlation of intensities | 2-3 | $[-1, 1]$ |

Table 2: Summary of available quantitative measures in FlowFI.

| Abbreviation | Description | Quantification |
| --- | --- | --- |
| SYCybyD | Symmetry of CD41 signal around the DRAQ5 centroid | Symmetry |
| CorrDC | Spatial correlation of DRAQ5 with cellular (forward scatter) | Spatial Corr. |
| CSD | Solidity (convexity) of the cell mask | Solidity |
| CySD | Solidity (convexity) of the FITC (cytoplasm) mask | Solidity |
| DSD | Solidity (convexity) of the DRAQ5 (DNA) mask | Solidity |
| CSZ | Cell size in forward scatter using pixel area | Area |
| ColCyC | The fraction of FITC signal contained inside the cell mask | Colocalisation |
| DSZ | DRAQ5 (DNA signal) mask size using pixel area | Area |
| ANGCybyD | Angular momentum of FITC signal around DRAQ5 | Angular Mom. |
| SPDCyC | Radial skewness of FITC centred around DRAQ5 | Relative Skew. |

Table 3: Summary of novel parameters used in MK purification and parameter refinement.
